## Supplemental Materials for "Conformational dynamics of a nicotinic receptor neurotransmitter site"

**Supplementary Materials**

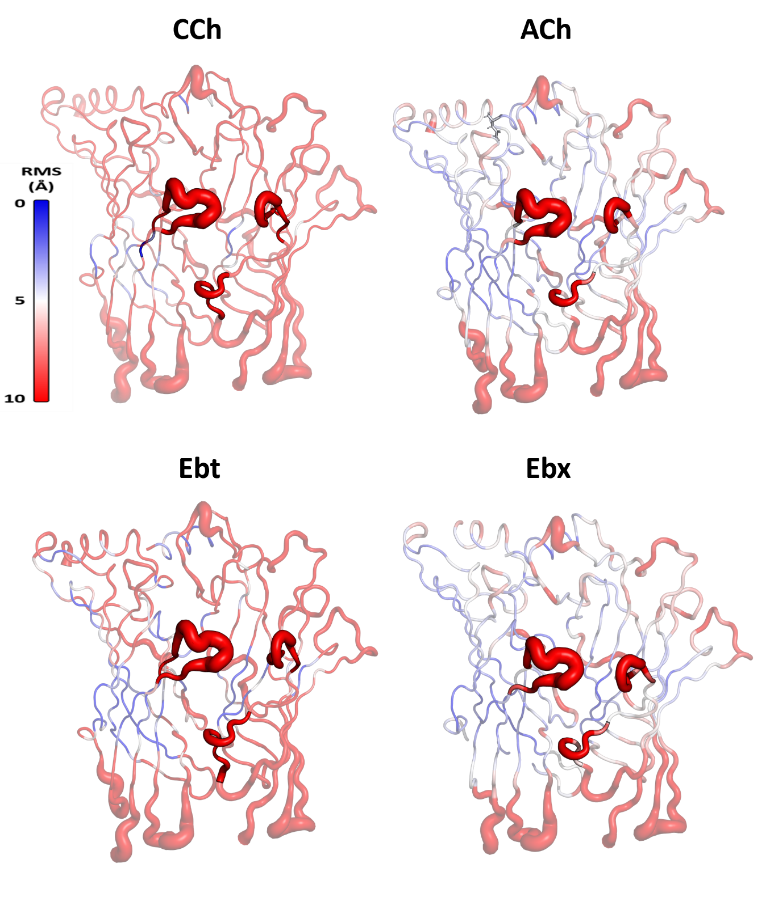

**Figure 2-figure supplement 1. Root Mean Square Fluctuation (RMSF) of C_α_ for extracellular domain (ECD) of α-δ subunits.** RMSF for carbamylcholine (CCh), acetylcholine (ACh), epibatidine (Ebt), and epiboxidine (Ebx) (structures in Figure 4A). Greatest fluctuations are in loops C and F that surround the orthosteric pocket; red-blue is greatest-least fluctuations.

RMSD (Å)

Time (ns)

System

Binding pocket

residues

Intra-facial

residues

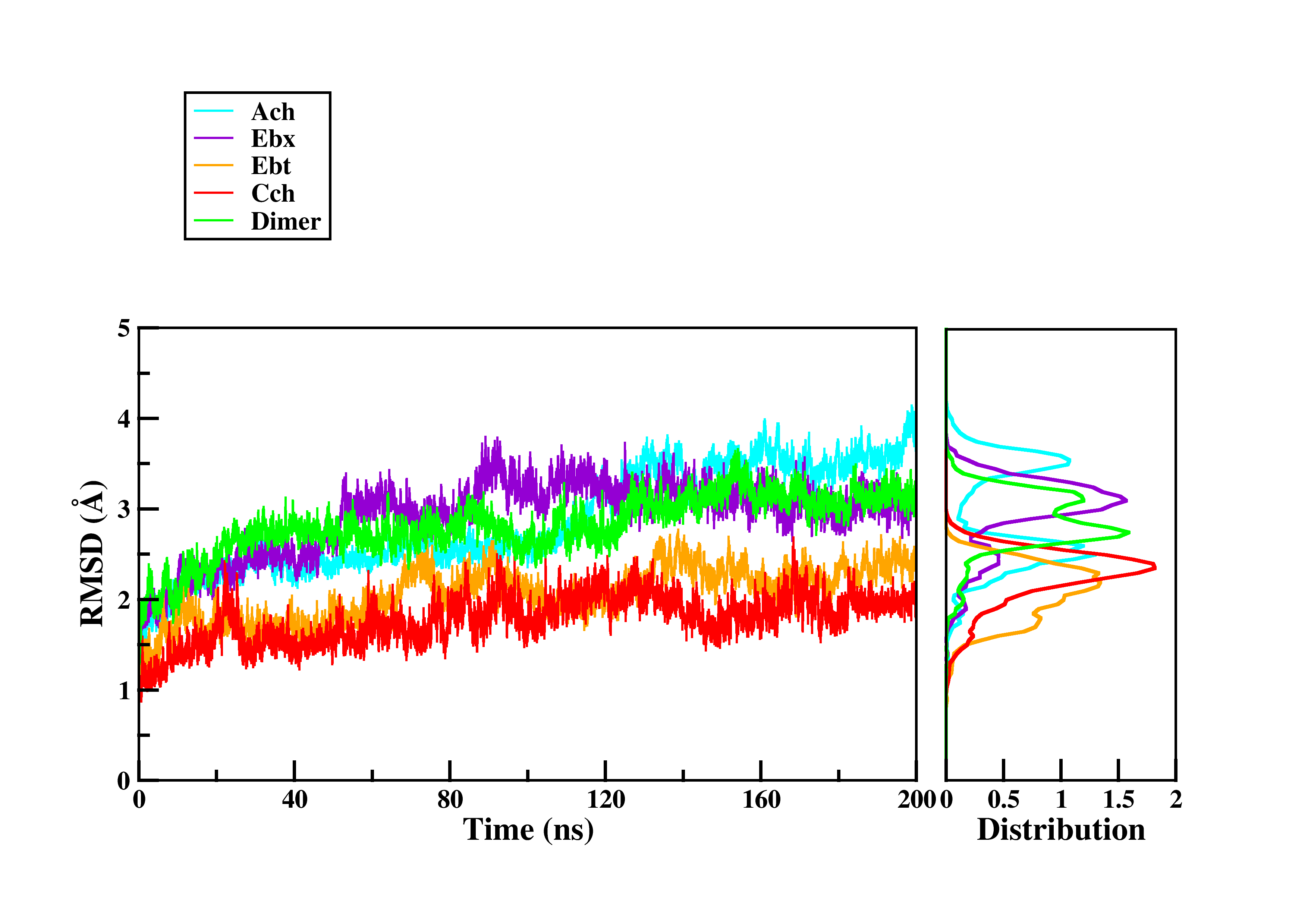

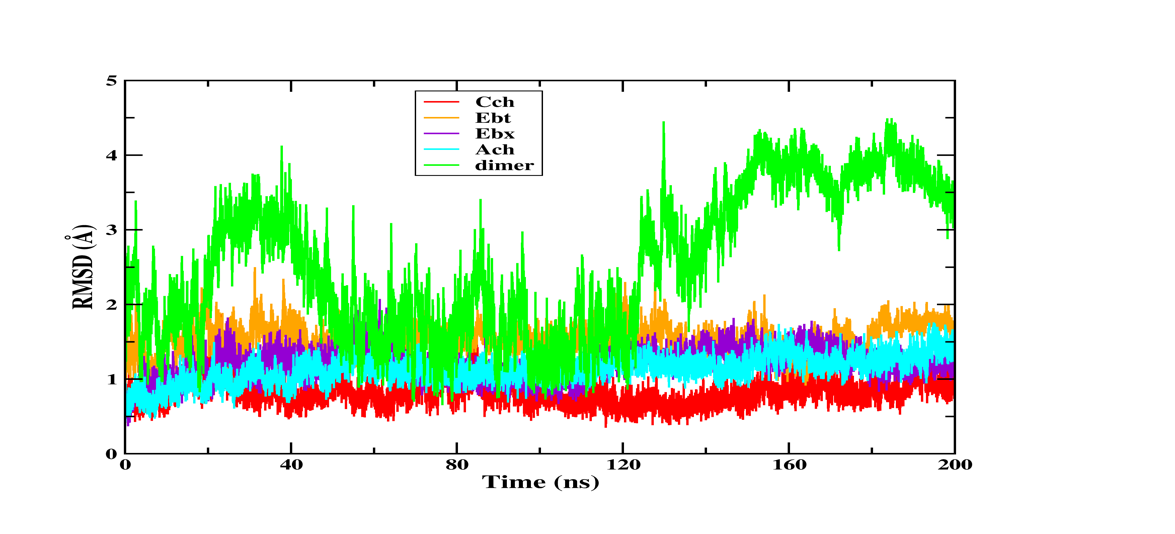

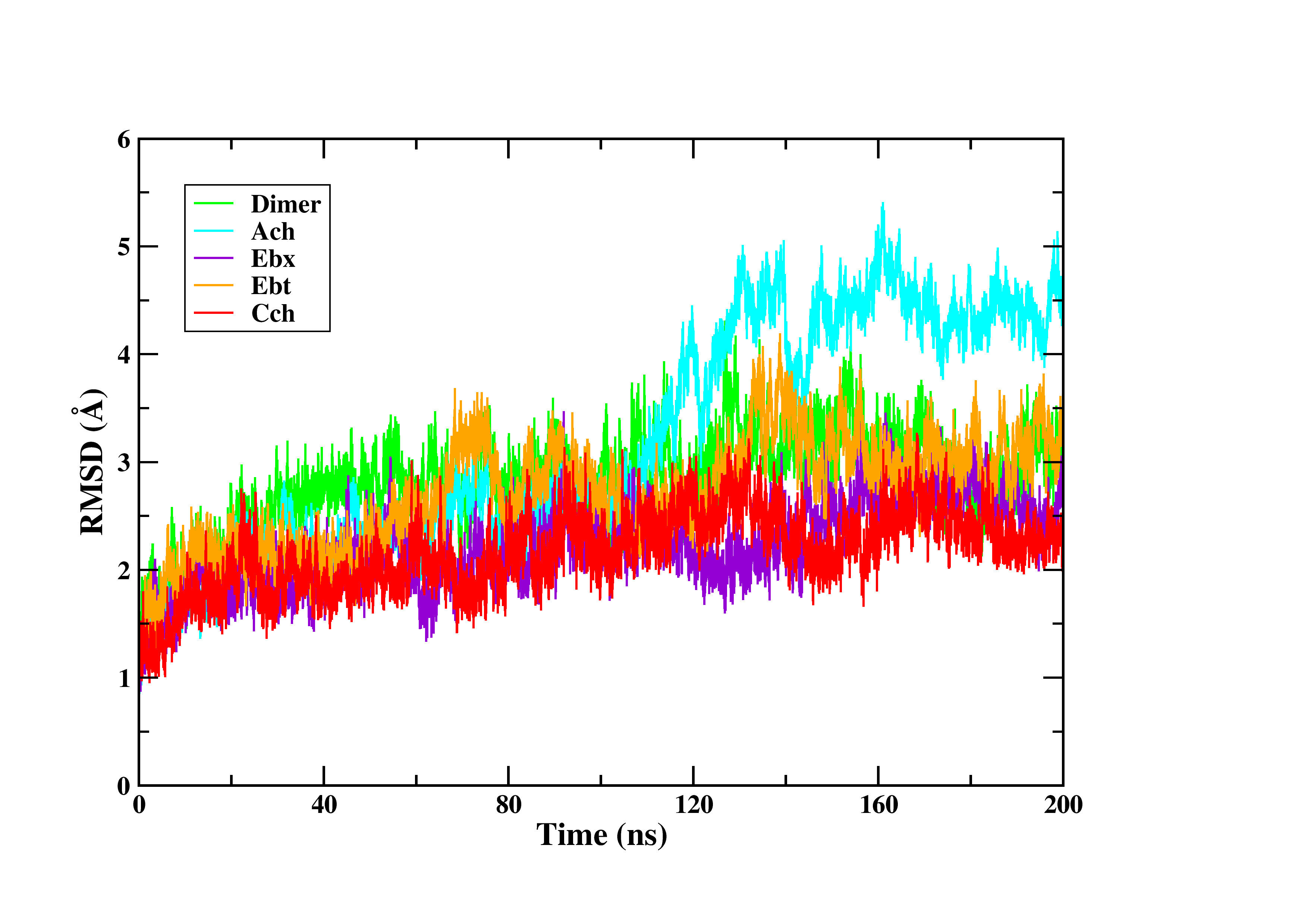

0

40

80

120

160

200

0

1

2

3

4

6

5

0

1

2

3

4

5

1

2

3

4

5

0

**Figure 2-figure supplement 2. RMSD of pocket residues during MD simulations.** The root-mean-square deviation (RMSD) of binding pocket residues relative to their initial positions over the course of 200 ns MD simulations for each agonist (CCh in red, EBt in orange, EBx in purple, ACh in blue and apo in green). Agonists stabilize pocket residues (middle) but intra-facial residues (near the TMD and adjacent subunits; lower) fluctuate similarly with and without agonists.

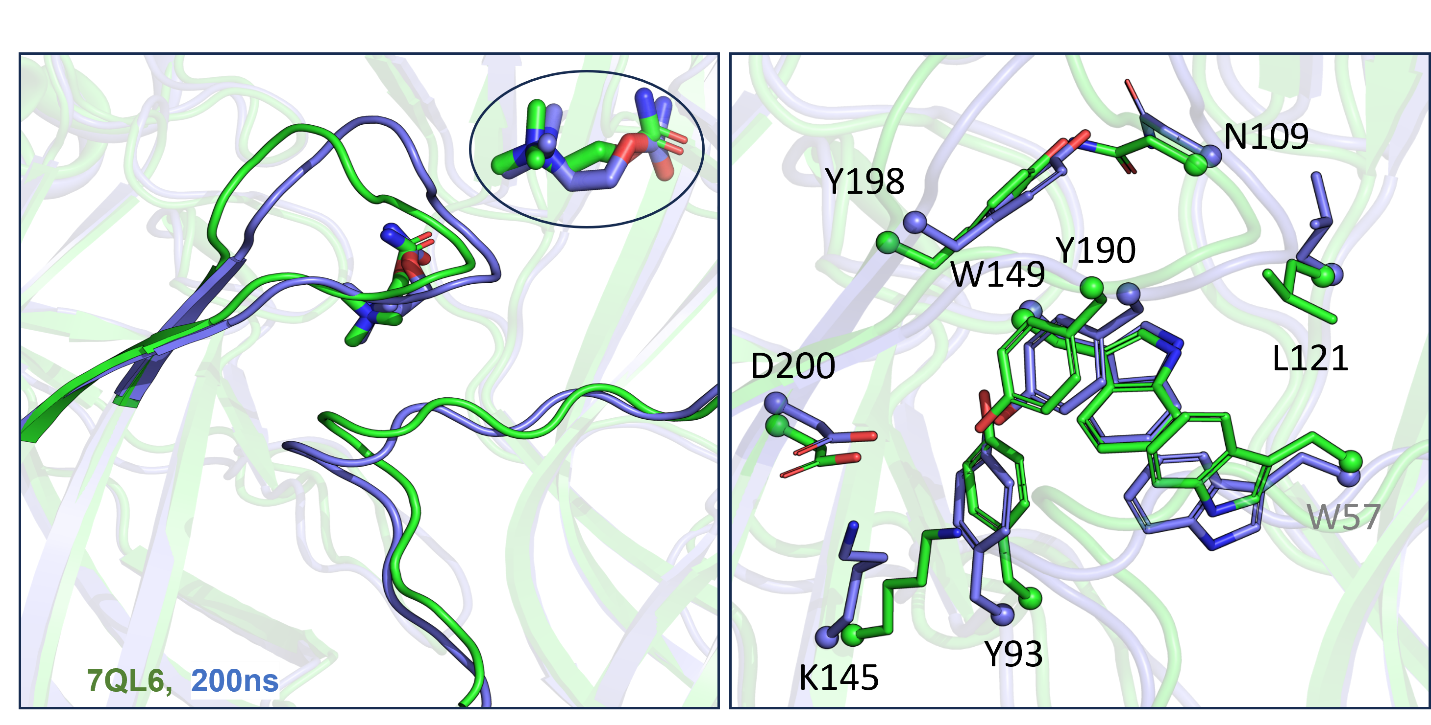

**Figure 2-figure supplement 3. Alignment of MD simulations (CCh).** Left, loops. An overlay of the final MD configuration with CCh (putative AC**_H_**; blue) and the cryo-EM desensitized (AD**_H_**) structure with CCh (7QL6.PDB, green) shows strong overlap in loops C (top left) and F (bottom right) (RMSD=0.3). Insert: the agonist (*trans* orientation) also overlaps. In experiments, AC**_H_** and AD**_H_** have the same CCh binding free energy. Right, side-chains. The final MD structure (blue) aligns approximately with the desensitized, high-affinity conformation of the pocket (green).

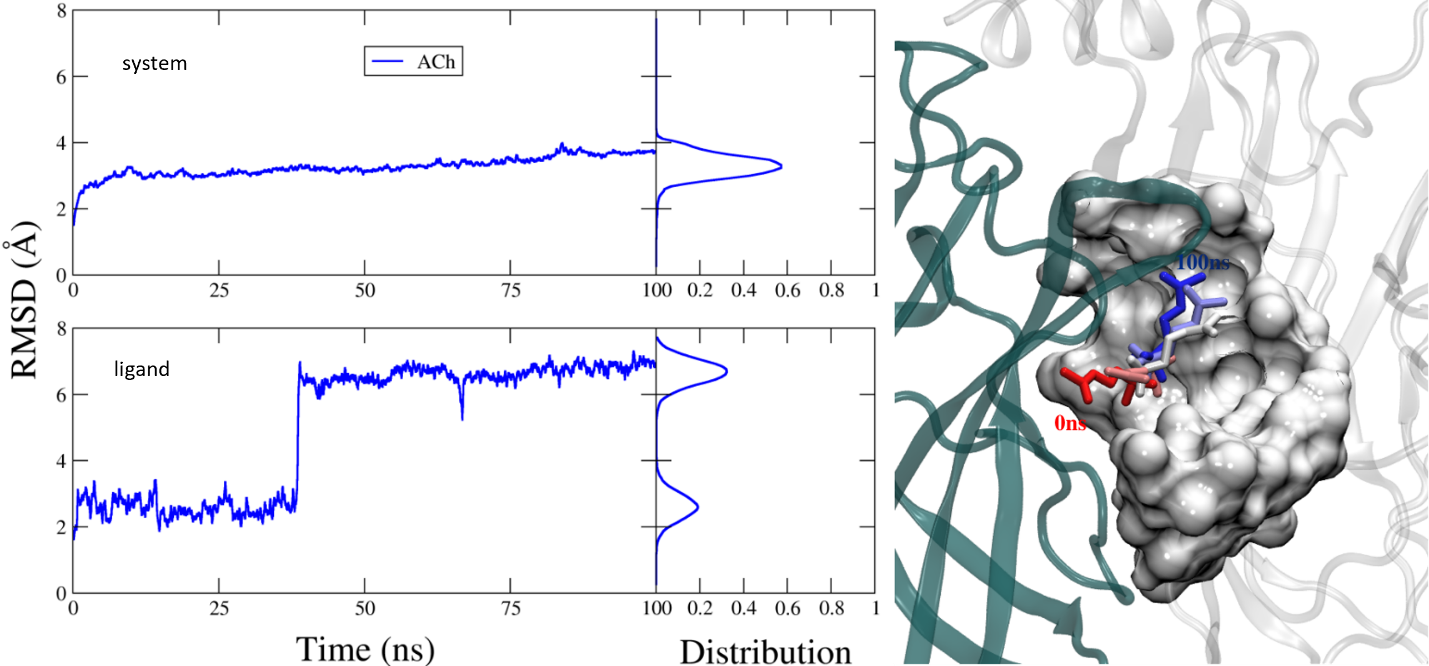

**Figure 2-figure supplement 4. MD simulation of the pentameric system.** Left, RMSD of the pentamer (top) and the ligand (bottom), in complex with docked acetylcholine (ACh) (100 ns MD simulation). Side panels are distribution plots. Right, the m1-m2-m3 changes in CCh conformation in the orthosteric cavity. The agonist's orientation flips from *cis* (m1, at 0 ns; red) to *trans* (m3, at 200 nS; blue). m2 is in between. Similar structural changes are apparent in the dimer.

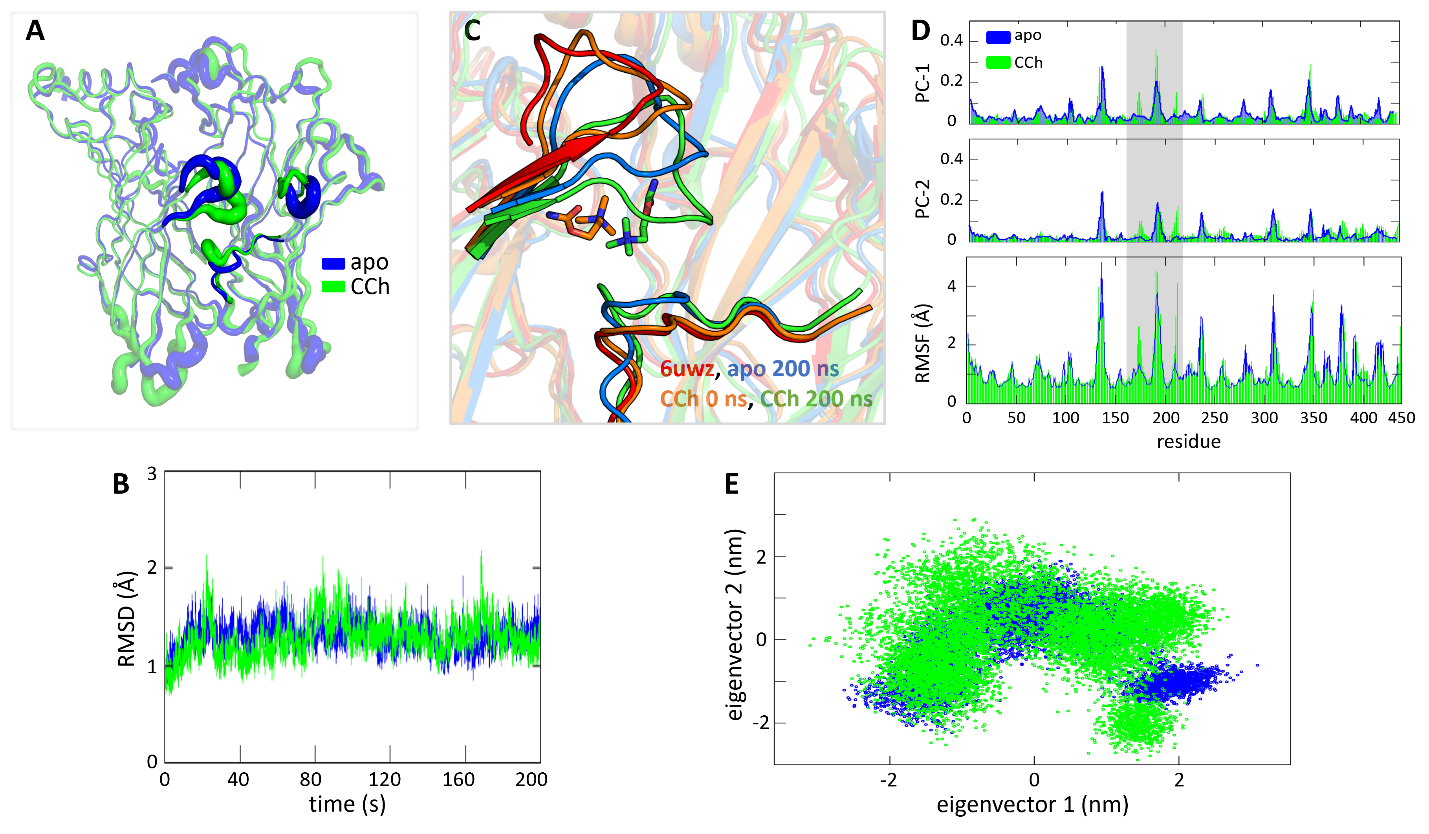

**Figure 2-figure supplement 5. Conformational convergence between apo (blue) and with CCh (green)**. A. RMSF of C_α_ ECD shows similar regions of fluctuations, loops C and F that surround the pocket. B. RMSD of the system C_α_. **C.** Closeup of the α−δ orthosteric pocket showing displacements of loop C (flop) and loop F; blue, apo; red, 6UWZ; orange CCh 0 ns; green, CCh 200 ns. D. RMSF shows loop C movement (grey highlight) is closely aligned with key conformational shifts observed in the primary two principal components (PC1 and PC2). **E.** A 2D scatter plot of the first two principal components derived from the simulation data for apo superimposed with CCh; cluster overlaps indicate that similar stable conformations were explored.

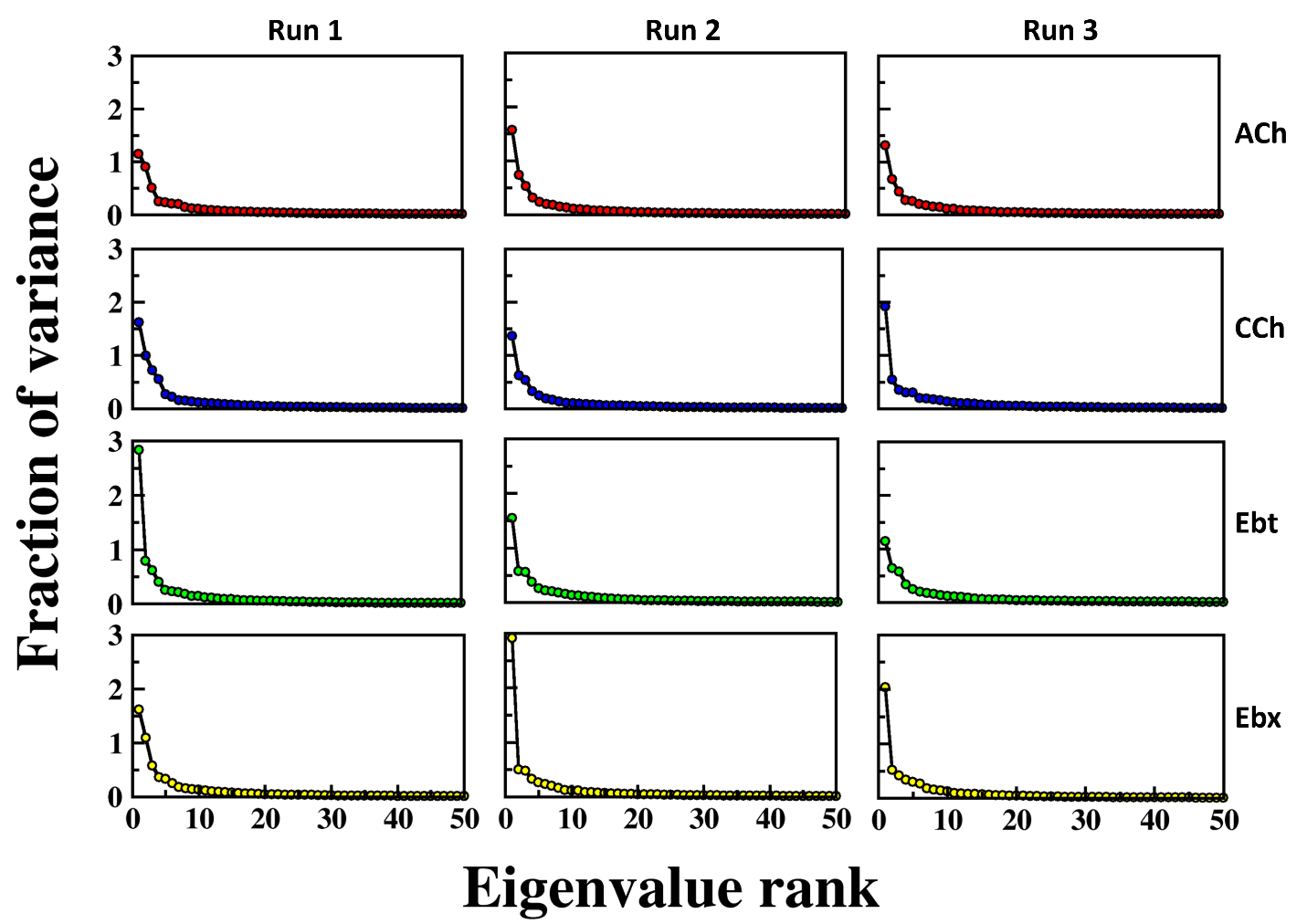

**Figure 3-figure supplement 1. Fraction of variance explained by eigenvalue rank for different simulation runs and conditions.** The plots show the fraction of variance (y-axis) as a function of eigenvalue rank (x-axis) for PCA. columns, 3 independent simulation runs (RUN1, RUN2, and RUN3); rows, 4 different conditions or datasets, indicated by different colors (red, blue, green, yellow). Each plot demonstrates that the first few principal components capture the majority of the variance, with a rapid decrease in the fraction of variance as eigenvalue rank increases. This pattern is consistent across all runs and conditions, indicating the robustness and reproducibility of the PCA results.

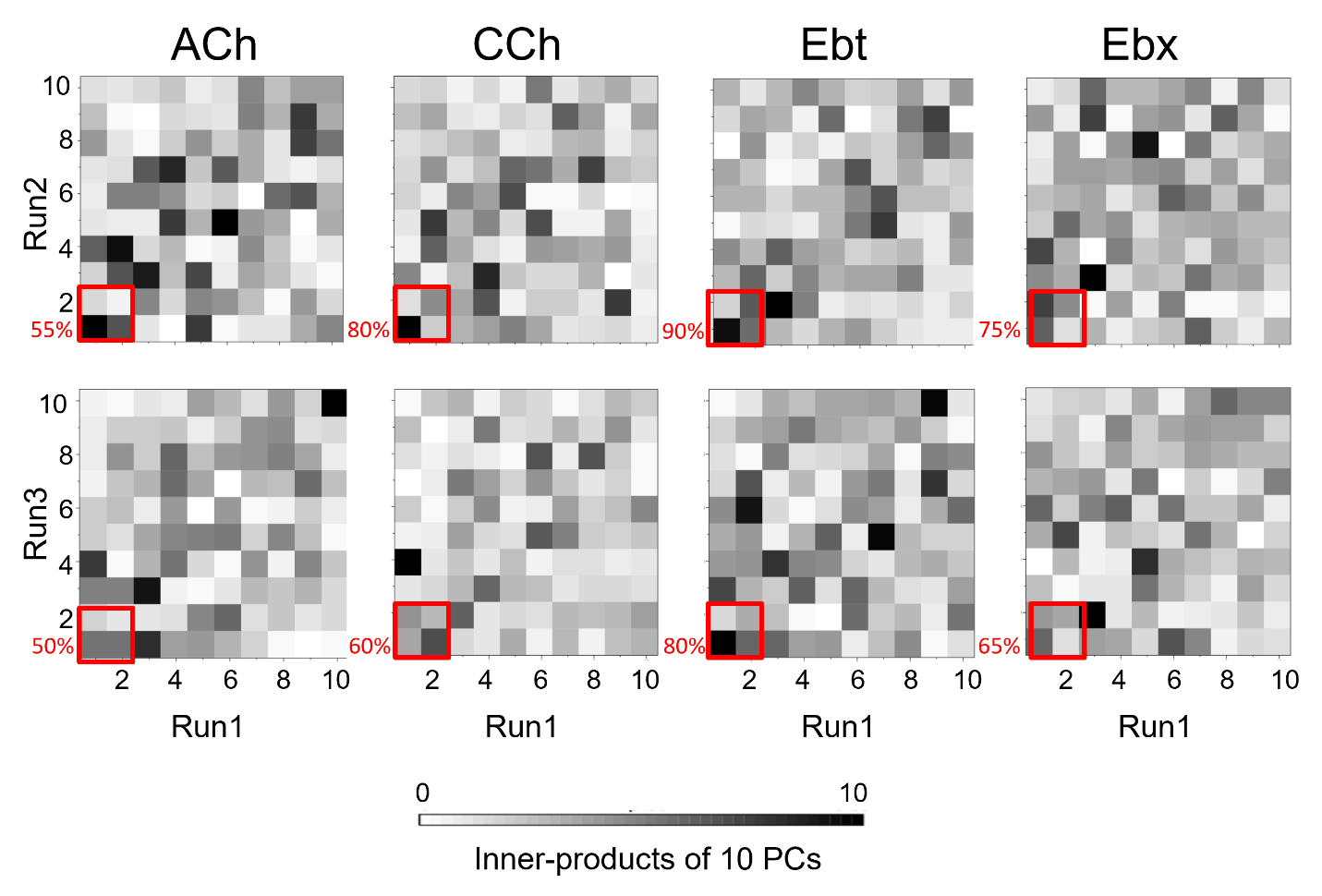

**Figure 3-figure supplement 2. Inner product heatmaps of principal components (PCs) from independent MD simulations:** The heatmaps display the inner product values of the top 10 PCs obtained from pairwise comparisons of three independent simulation runs for four systems - ACh, acetylcholine; CCh, carbamylcholine; Ebt, epibatidine and Ebx, epiboxidine. The color gradient from white to black indicates the magnitude of the inner product, with white representing an inner product of 0 (no correlation) and black representing an inner product of 10 (perfect correlation). The red boxes in the heatmaps correspond to PC-1 and PC-2 (percentage similarity between the runs indicated).

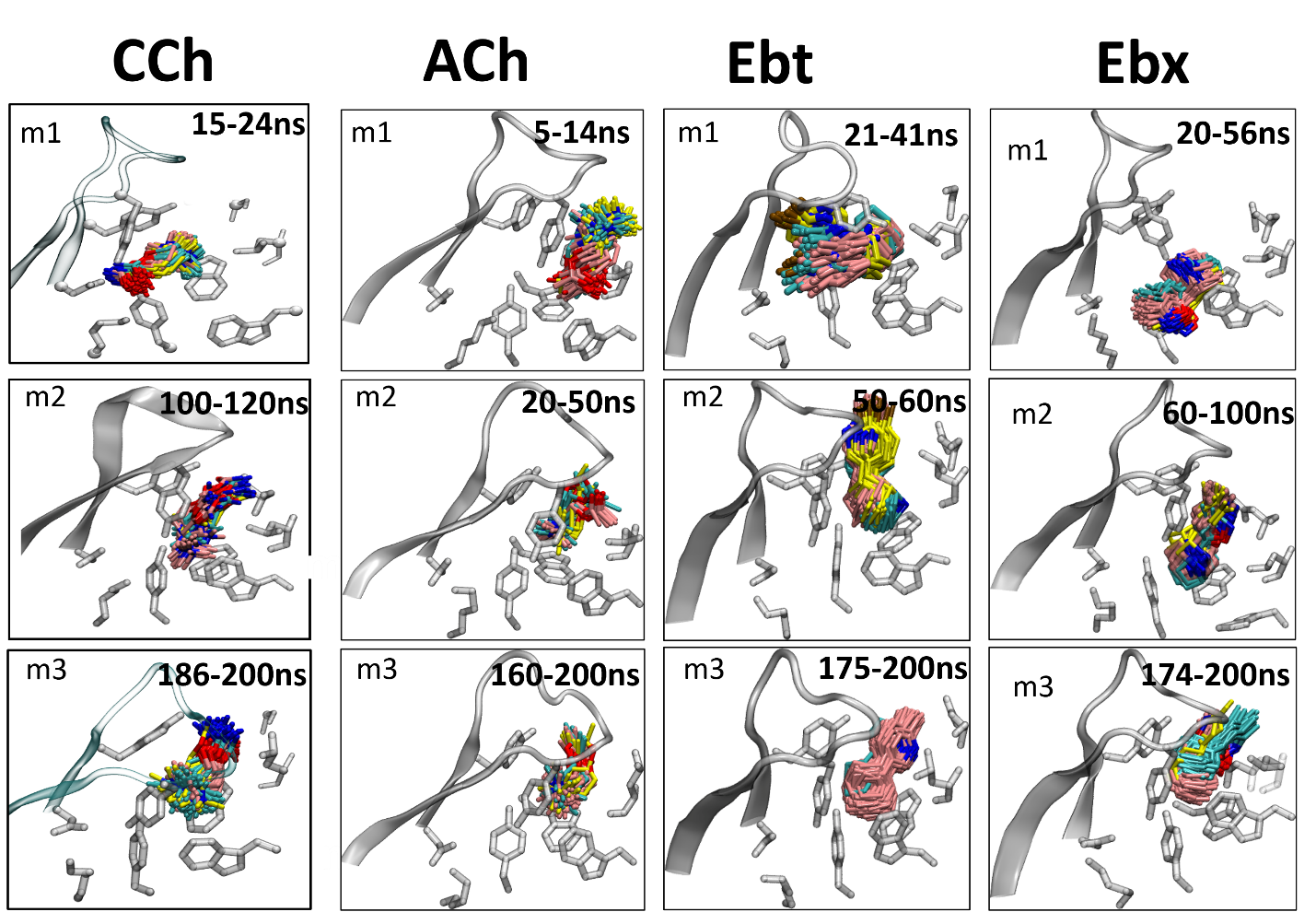

**Figure 3-figure supplement 3. Cluster analysis of ligand poses.** Three prominent poses of the ligand for each of the three energy minima (m1, m2, m3) were identified using cluster analysis in VMD. The top three clusters, with RMSD of ≤1Å each, display similar ligand orientations across the energy minima. The analysis was performed on frames extracted from the bottom of the wells apparent in the PCA. The temporal order in simulations is: m1 (start) < m2 (middle) < m3 (end). The agonists are carbamylcholine (CCh), acetylcholine (ACh), epibatidine (Ebt) and epiboxidine (Ebx). Three prominent clusters are colored by heteroatom with carbons in cyan, pink, and yellow respectively.

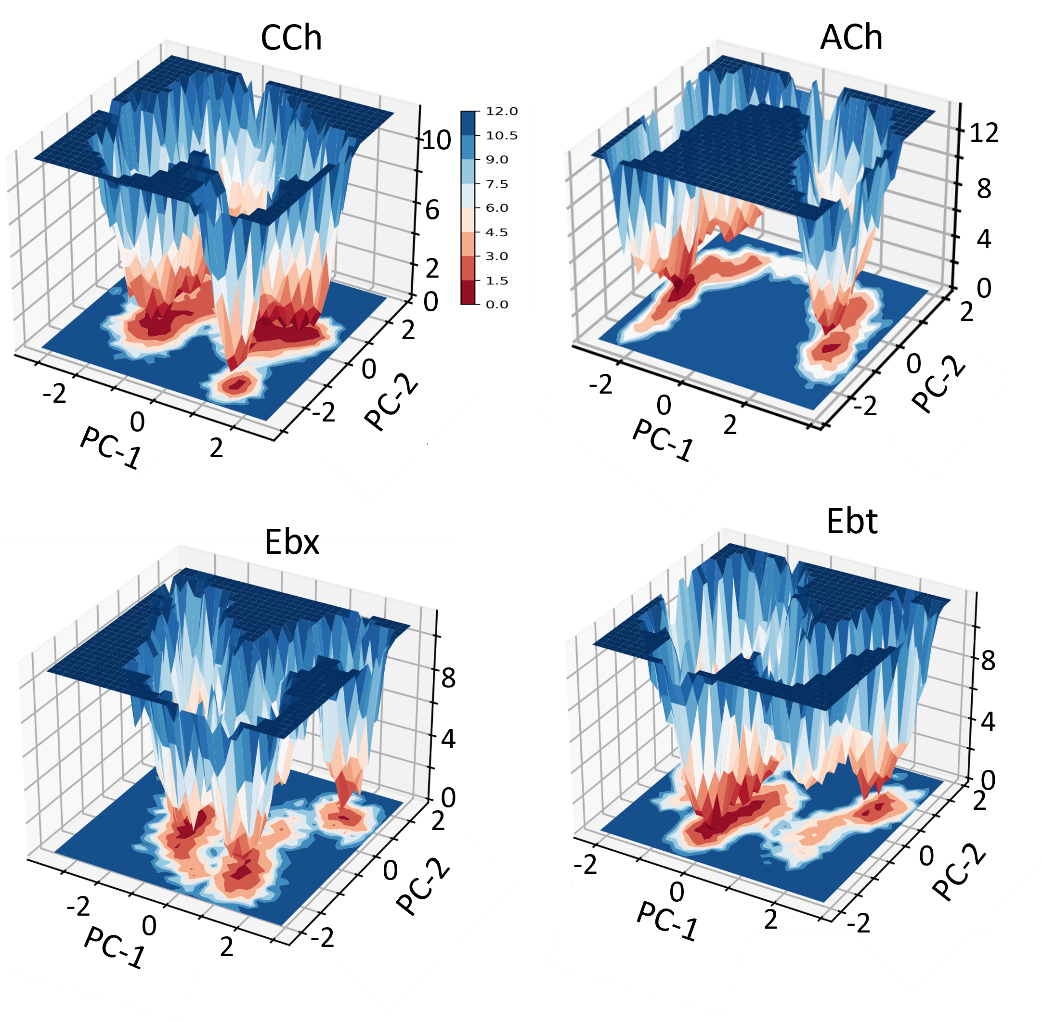

**Figure 3-figure supplement 4. Free energy landscapes as a function of PC1 and PC2.** The color scale represents the free energy values in kcal/mol; deeper wells (red) indicate more stable and shallower wells (blue) indicate less stable. All 4 agonists show 3 stable states (m1, m2, and m3). CCh, carbamylcholine; ACh, acetylcholine; Ebt, epibatidine and Ebx, epiboxidine.

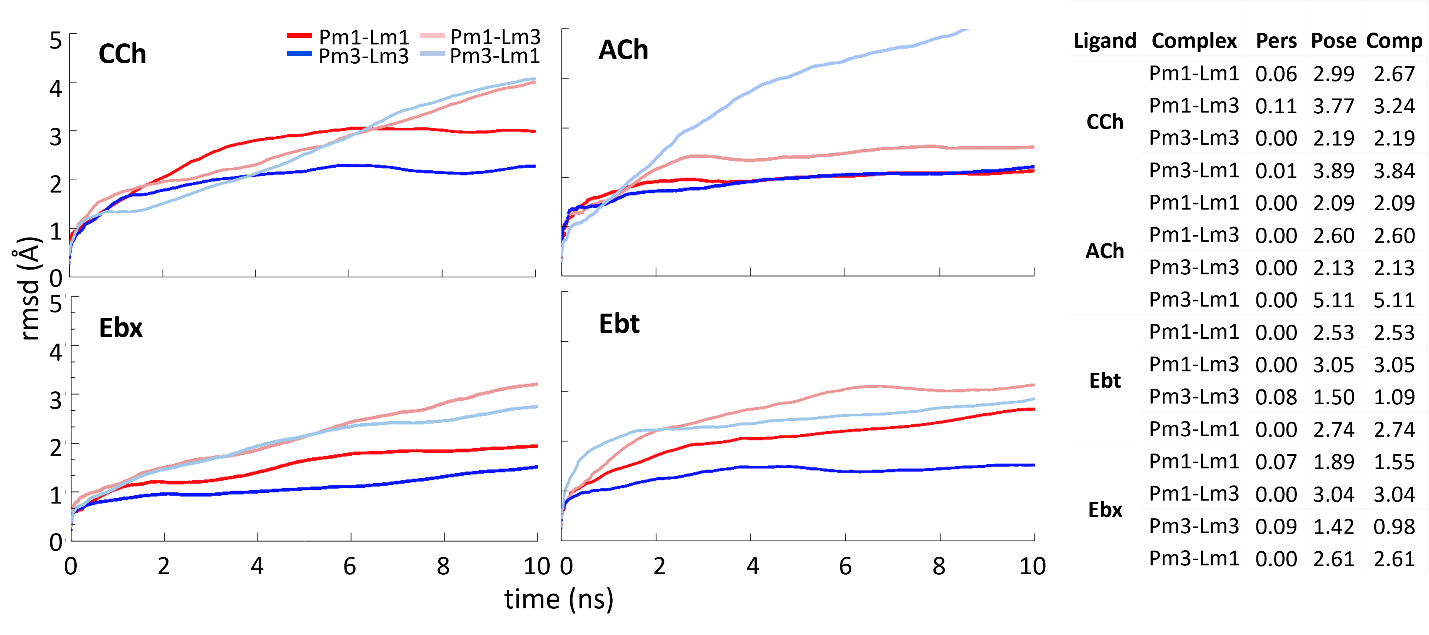

**Figure 3-figure supplement 5. Ligand-protein Complexes during binding pose metadynamics (BPMD).** An average RMSD plot of various ligand-protein complexes over the course of 10×10 ns metadynamics runs using PoseScore and PersScore derived values for four different agonists. The lines represent different ligand-protein combinations: Pm1-Lm1 (protein in m1 and ligand in m1, red), Pm1-Lm3 (protein in m1 and ligand in m3, light red), Pm3-Lm3 (protein in m3 and ligand in m3, blue), and Pm3-Lm1 (protein in m3 and ligand in m1, light blue). The plots indicate that the Pm3-Lm3 complexes show the least fluctuation, suggesting higher stability, while Pm1-Lm1 and the cross-docked complexes (Pm1-Lm3 and Pm3-Lm1) exhibit lower stability. Ttable, BPMD assesses the stability of binding poses using PoseScore and PersScore. PoseScore measures the RMSD of the ligand's heavy atoms relative to their initial coordinates, with lower values indicating higher stability. PersScore measures the persistence of important contacts, such as hydrogen bonds and pi-pi interactions, between the ligand and protein residues, with higher values indicating more stable complexes. The Composite Score (CompScore) combines PoseScore and PersScore (CompScore = PoseScore - 5 × PersScore) to provide an overall stability assessment, with lower CompScores indicating more stable ligand-protein complexes.

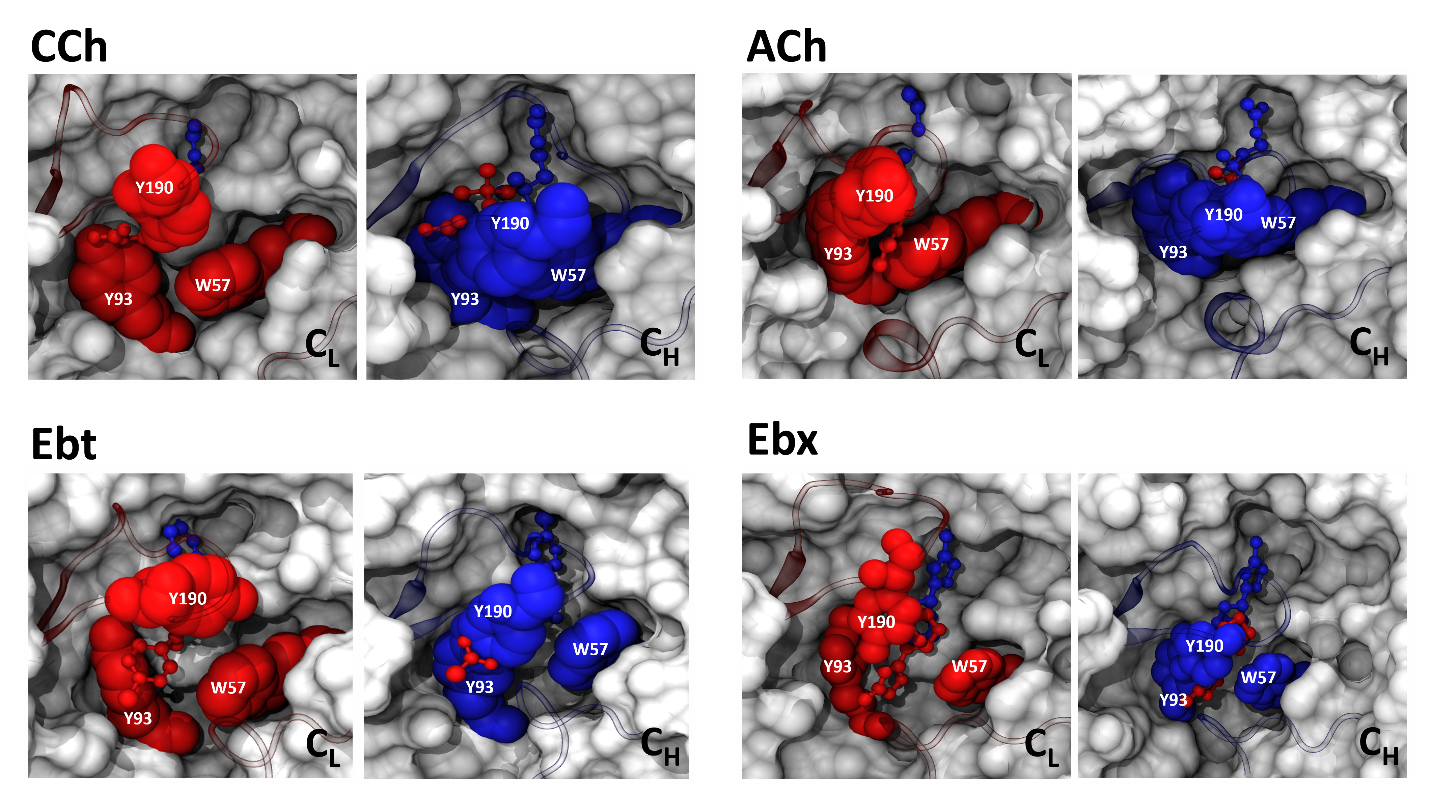

**Figure 5-figure supplement 1. Conformational change of the pocket cavity accommodates agonist re-orientation.** Red, low-affinity conformation (AC**_L_**); blue, high-affinity conformation (AC**_H_**). In AC**_L_** the ligand is *cis* and aromatic residues αY190, αY93, and δW57 are spread to accommodate the agonist's tail. In AC**_H_** the ligand is *trans* and αY190 is positioned to fill the αY93-δW57 gap, creating a unified surface that restrains the ligand's tail. Key residue sidechains are shown as spheres; agonist is ball-and-stick.

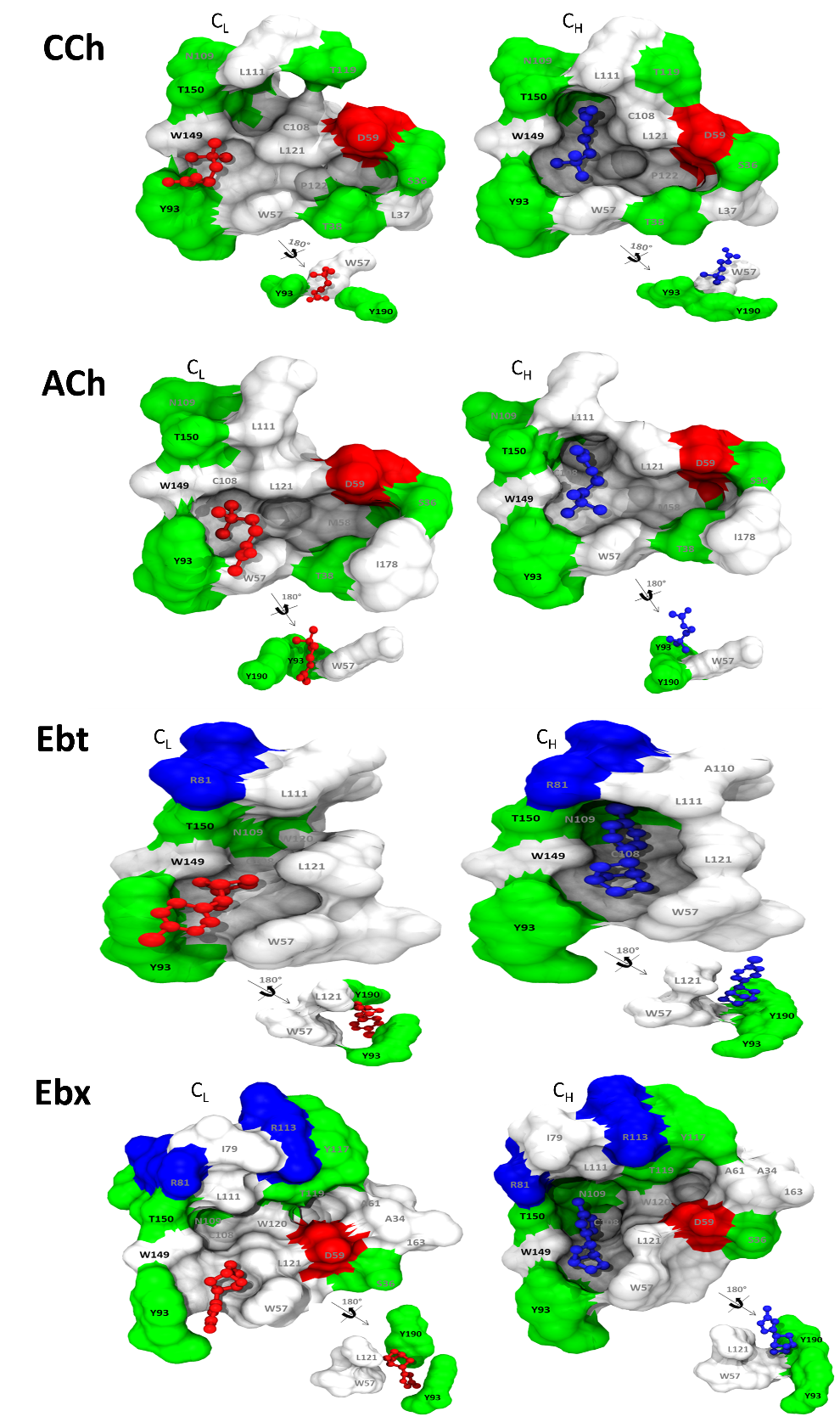

**Figure 6-figure supplement 1. Binding cavities (α−δ subunit interface)**. Left. Agonist (red) is low affinity (red: AC**_L_**, m1). Right, agonist is high affinity (blue; AC**_H_**, m3). All agonists (see Fig. S5) rotate about the cationic center (N^+^)-αW149 interface, so that the tail flips from being near αY93 in L (*cis*) to being in contact with the complementary-subunit surface in H (*trans*).

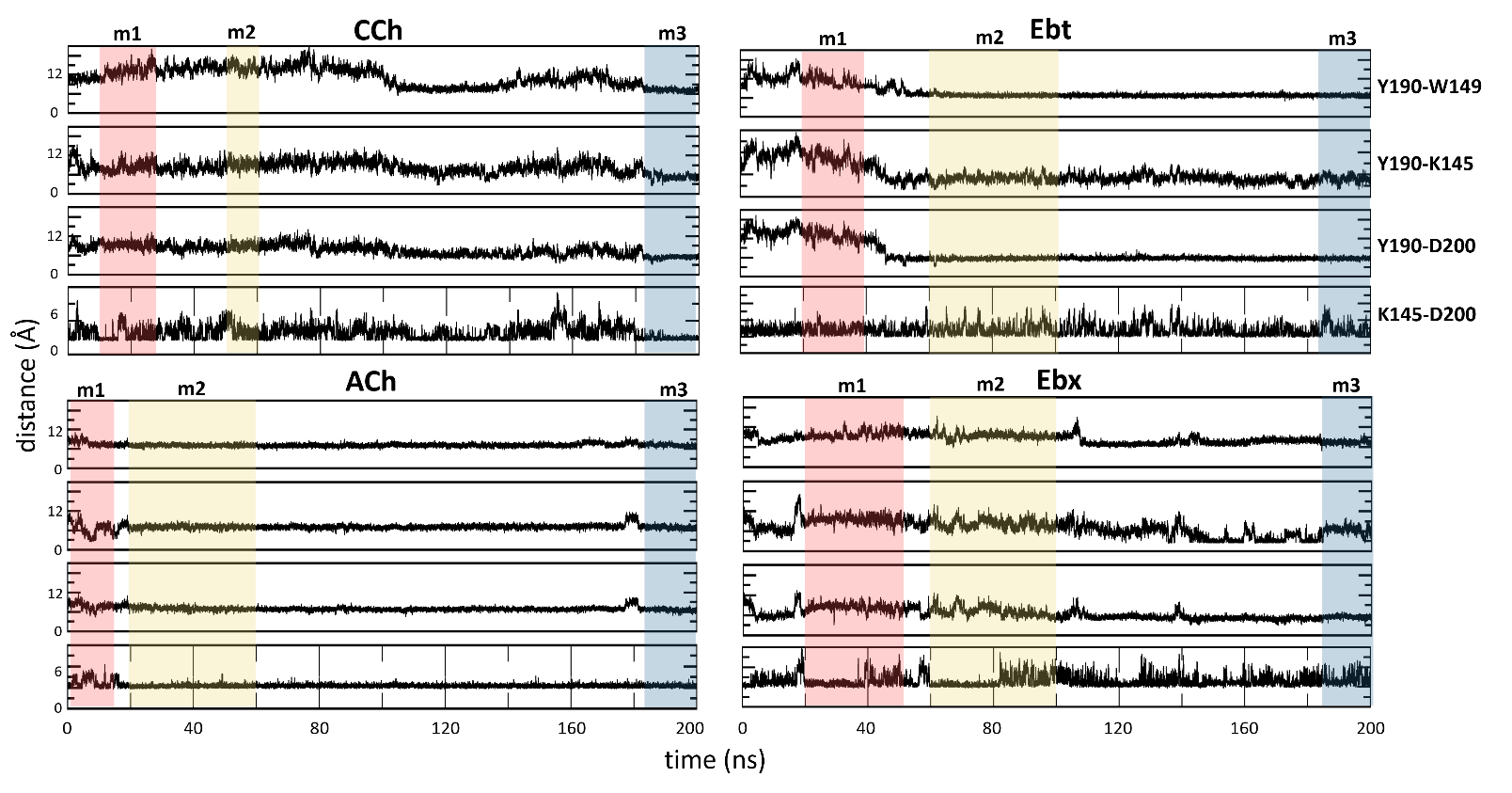

**Figure 6-figure supplement 2. Key residue distances over the course of MD simulation.** The distances in angstroms between specific functional groups of the hydroxyl group of αY190 to the indole of αW149, the amine of αK145, the carboxyl of αD200, and the salt-bridge distance between αK145 and αD200. The bottom energy minima for each system are highlighted: m1, red; m2, yellow; m3, blue.

|  | Time (ns) | System | RMSD (Å) | | | SD | SEM |
| --- | --- | --- | --- | --- | --- | --- | --- |
|  |  |  | Run1 | Run2 | Run3 |  |  |
| CCh | 200 | Complex | 1.84 | 2.20 | 2.19 | 0.21 | 0.12 |
|  |  | Ligand | 4.26 | 4.75 | 4.92 | 0.34 | 0.2 |
|  |  | Protein | 1.27 | 1.64 | 2.06 | 0.39 | 0.23 |
| ACh | 200 | Complex | 2.95 | 2.95 | 3.01 | 0.03 | 0.02 |
|  |  | Ligand | 6.83 | 6.83 | 5.57 | 0.73 | 0.42 |
|  |  | Protein | 2.43 | 2.44 | 2.53 | 0.05 | 0.03 |
| Ebt | 200 | Complex | 2.29 | 2.04 | 2.00 | 0.16 | 0.09 |
|  |  | Ligand | 4.37 | 3.85 | 3.08 | 0.65 | 0.37 |
|  |  | Protein | 1.14 | 1.58 | 1.55 | 0.25 | 0.14 |
| Ebx | 200 | Complex | 2.91 | 2.81 | 2.91 | 0.03 | 0.02 |
|  |  | Ligand | 5.46 | 5.59 | 5.46 | 0.73 | 0.42 |
|  |  | Protein | 2.90 | 2.22 | 2.90 | 0.05 | 0.03 |

**Figure 2-Source Data 1. MD simulation.** Average root-mean-square deviation (RMSD) values for protein, ligand, or the entire complex (protein + ligand) across individual MD simulation runs. SD, standard deviation; SEM, standard error of mean.

| **PCs** | **Cumulative contribution %** | | | | |
| --- | --- | --- | --- | --- | --- |
|  | **Apo** | **CCh** | **ACh** | **Ebt** | **Ebx** |
| PC1 | 20.72 | 31.98 | 49.55 | 23.25 | 30.14 |
| PC2 | 27.19 | 35.47 | 57.23 | 31.86 | 37.81 |
| PC3 | 38.49 | 45.24 | 62.15 | 40.29 | 44.01 |
| PC4 | 44.36 | 52.69 | 65.9 | 45.99 | 49.09 |
| PC5 | 48.58 | 56.38 | 69.18 | 49.93 | 53.52 |
| PC6 | 52.33 | 59.42 | 71.4 | 53.3 | 57.51 |
| PC7 | 55.67 | 61.58 | 73.06 | 56.39 | 60.17 |
| PC8 | 58.44 | 63.57 | 74.66 | 59.09 | 62.53 |
| PC9 | 60.78 | 65.39 | 76.09 | 61.45 | 64.61 |
| PC10 | 62.76 | 66.99 | 77.28 | 63.48 | 66.46 |

**Figure 3-Source Data 1. Cumulative contribution of principal components (PCs) to the variance in MD Simulations:** The table presents the percentage of total variance in the molecular motion that is accounted for by the first ten principal components in five different simulation conditions: Apo; CCh, carbamylcholine; ACh, acetylcholine; Ebt, epibatidine, and Ebx, epiboxidine. Each row corresponds to a principal component (PC1 through PC10), while the columns list the cumulative percentage of motion variance that the PCs account for in each simulation condition.

|  |  | **window (ns)** | **n frames** | **ΔH**  **(SD, SEM)** | **TΔS**  **(SD, SEM)** | **ΔG calculated** | **ΔG experimental** | **η calculated** | **η**  **experimental** |
| --- | --- | --- | --- | --- | --- | --- | --- | --- | --- |
| **CCh** | **m1** | 15-24 | 226 | -27.61  (7.66, 0.49) | -22.88  (5.82, 2.37) | -4.73 | -4.44 | 0.52 | 0.52 |
|  | **m2** | 100-120 | 273 | -24.41  (5.68, 0.79) | -17.54  (3.49, 1.42) | -6.87 |  |  |  |
|  | **m3** | 186-200 | 207 | -26.18  (5.75, 0.47) | -16.25  (7.76, 2.24) | -9.93 | -9.20 |  |  |
| **ACh** | **m1** | 5-14 | 100 | -20.83  (8.11, 1.13) | -14.27  (6.51, 3.25) | -6.56 | -5.11 | 0.47 | 0.50 |
|  | **m2** | 20-50 | 359 | -27.92  (3.01, 0.29) | -16.72  (2.53, 1.03) | -11.2 |  |  |  |
|  | **m3** | 160-200 | 266 | -27.00  (2.88, 0.87) | -14.57  (6.28, 3.63) | -12.43 | -10.31 |  |  |
| **Ebt** | **m1** | 21-41 | 591 | -27.64  (3.83, 0.27) | -13.42  (4.70, 1.42) | -14.22 | -6.20 | 0.42 | 0.42 |
|  | **m2** | 50-60 | 325 | -24.21  (4.25, 0.42) | -13.50  (6.81, 2.05) | -10.71 |  |  |  |
|  | **m3** | 175-200 | 996 | -41.77  (3.99, 0.28) | -17.23  (6.86, 2.07) | -24.54 | -10.60 |  |  |
| **Ebx** | **m1** | 20-56 | 1284 | -20.51  (2.98, 0.46) | -12.85  (5.49, 1.66) | -7.66 | -5.40 | 0.48 | 0.46 |
|  | **m2** | 60-100 | 504 | -21.55  (3.56, 0.28) | -14.39  (10.46, 3.15) | -7.16 |  |  |  |
|  | **m3** | 174-200 | 562 | -33.95  (2.92, 0.88) | -19.33  (4.39, 1.32) | -14.62 | -9.96 |  |  |

**Figure 4-Source Data 1. Table of calculated and experimental binding energies (kcal/mol):** Agonists, see Fig. 2A. *In silico* (calculated) values were from frames from each PCA population (m1, m2 and m3; Fig. 3) after cluster analysis of ligand orientation. *In vitro* (experimental) values were from electrophysiology measurements. window, frame times for each population; n frames, number of simulation frames from three prominent clusters chosen for MM-PBSA analysis; ΔH, change in binding enthalpy; ΔS change in binding entropy (T, absolute temperature); ΔH, change in Gibbs free energy (ΔG=ΔH-TΔS); η, efficiency (1-ΔG_LA_/ΔG_HA_). SD, standard deviation; SEM, standard error of mean. Data are shown as bar graphs in Figure 4.

|  |  | EEL | Vdw | EPB | ENPOLAR | ΔG gas | ΔG solv |
| --- | --- | --- | --- | --- | --- | --- | --- |
| CCh | m1 | -333.79 (11.75, 0.76) | -21.83  (2.63, 0.17) | 330.28 (12.61, 0.81) | -2.17  (0.08, 0.01) | -355.72 (11.88, 0.76) | 328.10  (12.57, 0.81) |
|  | m2 | -320.28 (11.95, 1.67) | -25.53  (3.28, 0.46) | 323.63 (11.34, 1.58) | -2.23  (0.12, 0.01) | -345.81 (13.66, 1.91) | 321.40  (11.36, 1.59) |
|  | m3 | -328.51  (8.58, 0.69) | -30.64  (2.12, 0.17) | 335.07  (7.34, 0.59) | -2.08  (0.08, 0.01) | -359.15  (9.23, 0.74) | 332.98  (7.35, 0.59) |
| ACh | m1 | -304.06 (11.36, 3.88) | -22.44  (5.56, 0.78) | 305.64 (12.44, 3.75) | -2.21  (0.06, 0.02) | -324.26 (17.16, 5.17) | 303.43  (12.43, 3.74) |
|  | m2 | -311.50  (7.67, 0.76) | -27.94  (1.88, 0.19) | 313.74  (6.63, 0.66) | -2.23  (0.07, 0.01) | -339.43  (7.89, 0.78) | 311.51  (6.65, 0.66) |
|  | m3 | -292.97  (5.14, 1.55) | -29.15  (1.96, 0.59) | 297.33  (4.66, 1.40) | -2.23  (0.05, 0.01) | -322.10  (5.71, 1.72) | 295.10  (4.67, 1.4101) |
| Ebt | m1 | -354.29 (13.62, 0.96) | -28.18  (2.13, 0.15) | 357.44 (13.74, 0.96) | -2.62  (0.07, 0.01) | -382.46 (13.82, 0.97) | 354.82  (13.73, 0.9691) |
|  | m2 | -349.73 (10.40, 1.03) | -29.63  (2.54, 0.25) | 357.78  (9.80, 0.97) | -2.62  (0.08, 0.01) | -379.36 (10.46, 1.04) | 355.15  (9.80, 0.97) |
|  | m3 | -359.83  (7.83, 0.55) | -37.95  (1.95, 0.14) | 358.39  (7.13, 0.50) | -2.38  (0.04, 0.00) | -397.77  (7.74, 0.54) | 356.01  (7.13, 0.50) |
| Ebx | m1 | -319.00  (6.39, 0.99) | -27.13  (2.14, 0.33) | 328.16  (7.44, 1.16) | -2.55  (0.07, 0.01) | -346.13  (6.83, 1.06) | 325.61  (7.46, 1.16) |
|  | m2 | -321.95 (15.34, 1.21) | -23.34  (2.34, 0.18) | 326.33 (14.61, 1.15) | -2.61  (0.10, 0.01) | -345.26 (16.18, 1.27) | 323.71  (14.62, 1.15) |
|  | m3 | -321.11  (8.45, 2.55) | -30.67  (1.69, 0.51) | 320.25  (8.43, 2.54) | -2.43  (0.06, 0.01) | -351.77  (9.18, 2.76) | 317.81  (8.43, 2.54) |

**Figure 4-Source Data 2. MM-PBSA components of the free energy calculation (see Figure 4). EEL, electrostatic; VdW, van der Waals; EPB, electrostatic Poisson-Boltzmann; ENPOLAR, nonpolar solvation energy; ΔG gas, gas-phase free energy; ΔG solv, solvation free energy; SD, standard deviation; SEM, standard error of mean.**

|  |  | αK145-αD200 | αK145-αY190 | αD200-αY190 | N^+^-αW149 | loop C (αC192) |
| --- | --- | --- | --- | --- | --- | --- |
| CCh | m1 | 6.1 | 9.3 | 7.8 | 5.4 | 2.7 |
|  | m2 | 6.3 | 9.5 | 7.9 | 4.5 | 3.8 |
|  | m3 | 2.6 | 6.1 | 4.7 | 4.2 | 12.1 |
| ACh | m1 | 5.1 | 7.8 | 6.3 | 4.6 | 3.7 |
|  | m2 | 3.1 | 7.0 | 7.6 | 4.8 | 7.4 |
|  | m3 | 2.8 | 5.3 | 4.4 | 4.4 | 9.6 |
| Epi | m1 | 3.4 | 14.7 | 14.0 | 5.1 | 6.8 |
|  | m2 | 3.7 | 5.8 | 6.2 | 5.0 | 7.5 |
|  | m3 | 4.4 | 6.7 | 6.1 | 3.3 | 7.9 |
| Ebx | m1 | 4.4 | 8.5 | 7.8 | 6.3 | 5.5 |
|  | m2 | 3.8 | 9.7 | 8.3 | 5.4 | 4.7 |
|  | m3 | 5.4 | 8.0 | 5.2 | 3.8 | 9.7 |

**Figure 6-table supplement 1. Residue distances.** Distance measurements (Å) between functional groups. αK145 is to companion polar group; N^+^ is to center of indole ring; loop C is C_α_ of αC192 compared apo; m1, m2 and m3 structures taken from plateau regions of the trajectories.
